# Simultaneous Cancer Treatment with Photothermal Therapy and Chemotherapy using Gold nanorods coated with Methotrexate conjugated Hyaluronic acid

**DOI:** 10.64898/2026.04.16.719030

**Authors:** Nutan Shukla, Ratnesh Das

## Abstract

Upon near-infrared (NIR) irradiation, combined treatment comprising of photothermal therapy (PTT) and chemotherapy (CHT) offers synergistic effects by inducing localized heat to intended tumor sites and simultaneously allowed delivering drugs thus to minimize undesired side-effects but enhance cytotoxic therapies. In this study we developed a novel platform that enables simultaneously to respond light stimuli with localized heat and released drugs using drug contained gold nanorods (GNRs). Methotrexate (MTX), a model anticancer drug is attached through hydrolytic ester bonding to targeting molecular hyaluronic acid (HA) that is coated onto GNRs. Based on the rationale, HA provides a good scaffold for high biocompatibility to shield risky GNRs, targeting for a CD44 receptor, and easy chemical binding of drugs. Upon a single light irradiation, MTX-HA functionalized GNRs (MTX-HA @GNRs) provide localized heat to cancer areas for PTT and the elevated temperature accelerates hydrolytic cleavage of the ester bond onto GNRs in physiological condition for CHT, ultimately releasing MTX to cells. In contrast to previous combination therapies that do not concurrently offer heat and drugs upon light stimuli, our NIR triggered CHT with PTT provides clinically effective options with combinatorial treatment that possesses high efficacy resulted in *in vitro* tests.

Scheme 1.
Schematic illustration of our nanoplatform a) The light responsive combinational therapy using GNRs scaffold for photo thermal therapy and chemotherapy b) The formation of TGNRs@RHO.B-HA and their light responsive mechanism with active CD44 receptor binding affinity c) light triggered hydrolytic release of model drug Rhodamine.B from TGNRs@RHO.B-HA.

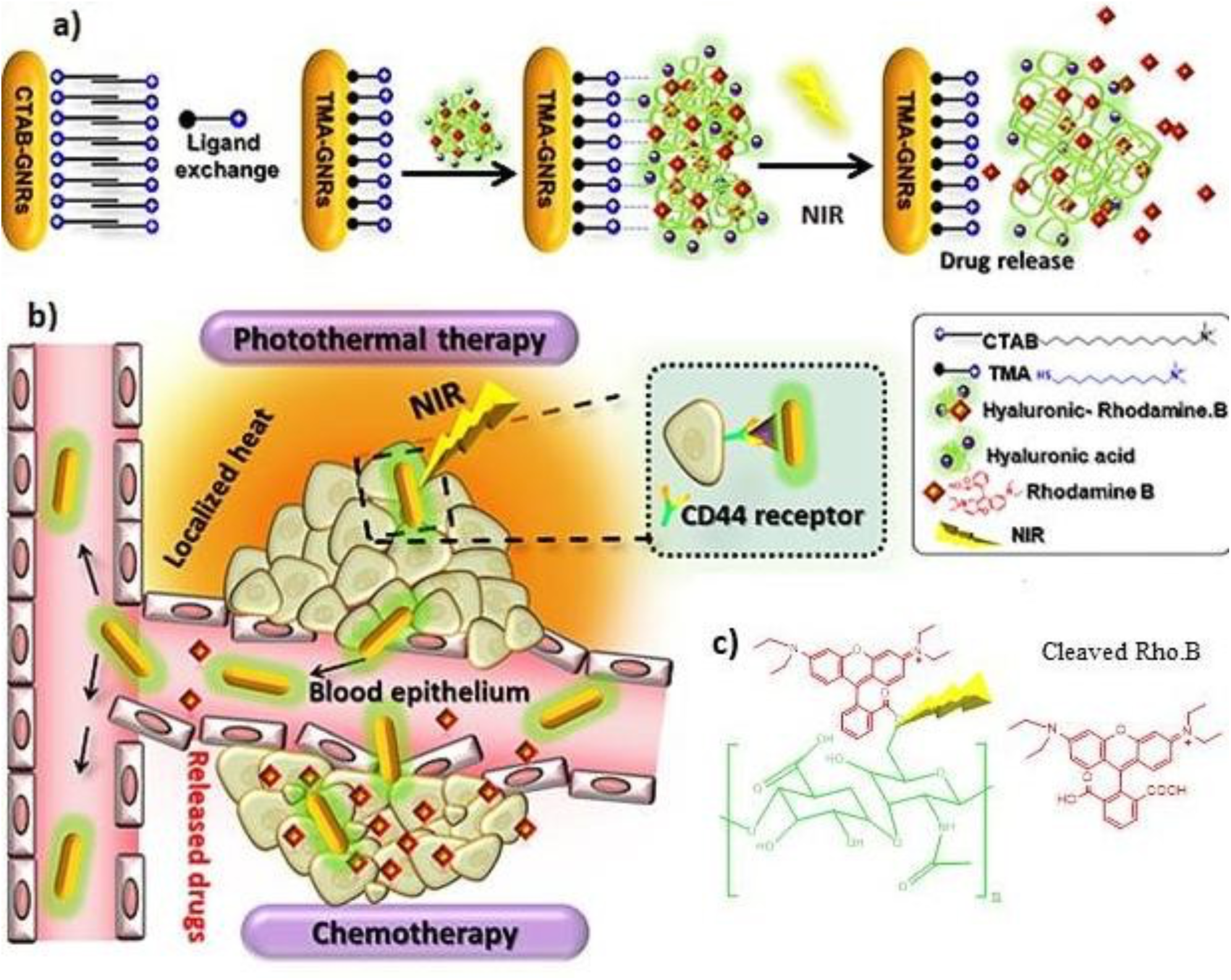

## Introduction

Cancer is characterized as malfunctioning of gene that results into uncontrolled proliferation of cells. **^[1, 2]^** According to the world health organization it is estimated one of the leading causes of mortality increasing every year. **^[3]^** Current treatment modalities such as surgery, irradiation therapy, chemotherapy, and immune-therapy have become therapy regimen. **^[4]^**Amongst, chemotherapy (CHT) the long-established form of treatment is widely applied due to high therapeutic efficacy. **^[5]^**Chemo-drug effectively restricts proliferation of cells by interfering their enzyme function and molecular mechanism. Further, it actively participates in tumor suppression, anti-angiogenesis, tumor quenching, and metastasis inhibition. **^[6]^**However, owing to some disadvantages such as non-specific delivery, high dosage requirement, and patient discomfort CHT frequently fails to maintain its therapeutic protocol. **^[7]^**To overcome these limitations of CHT, various theranostics are continuously being researched and combined to improve therapeutic effects. **^[8]^**In particular, to compensate for the lack of CHT, combinational therapy such as chemo-immunotherapy, chemo-radiotherapy, chemo-photodynamic therapy and chemo-photothermal therapy have become new paradigm. Amongst, the combination of CHT and PTT is one of the promising approaches to overcome systemic side effects by improving the therapeutic effect. **^[9]^**

NIR with the deep tissue penetration and low light toxicity is promising as trigger in drug delivery systems. **^[10]^**Hence, NIR responsive nanomaterial’s such as gold nanorods, gold nanocages, gold nanoshells, gold nanoparticle, graphene oxide, and carbon nanotubes have attracted attention in the oncology field as drug delivery system. **^[11]^** Especially, GNRs is preferred as smart material because of their multivalent properties like localized surface plasmon resonance (SPR), pharmacokinetics, optical property (light scattering), easy surface modification, antitumor efficacy and passive targeting ability. **^[12]^**The striking property of GNRs such as light conversion to localized cytotoxic heat allows programmable drug release on responding to NIR.**^[13]^** The high metabolic activity and heat sensitivity of microenvironment precisely elevates the localized temperature to 40^0^C-46^0^C upon NIR irradiation accelerated by GNRs. **^[20, 21]^** this leads to decreased hydrokinetic pressure which enhances tissue permeability, cell adhesion and vasodilation of blood capillaries. **^[14–17]^** the enhanced cell adhesion improves extravasation of chemo-drug at tumor site. **^[18, 19]^**

To overcome these obstacles and enhanced therapeutic efficacy Yang’s research group reported combinational effect of anticancer drug mediated via NIR responsive GNRs. **^[22]^** their study aimed to develop nanomedicine that constituted covalent modification of GNRs surface by camptothecin (CPT). **^[23]^** Further, on addition of NIR controlled release of drug was observed.

Unfortunately, its biocompatibility and active targeting effect was not reasonably concluded. **^[24]^**Considering above limitation we rationale that encapsulation of anticancer drug molecule into bioactive molecules of tumor microenvironment namely hyaluronic acid (HA), lipids, polymers, liposomes, polysaccharide preferentially enhances cellular uptake and biocompatibility. **^[25]^**These moieties actively recognize surface markers on cancer cells and enables specific penetration of chemo-drugs. **^[26]^**

Amongst, hyaluronic acid (HA) the natural polymer has been widely investigated and reported in many kinds of research.**^[27]^**Besides biological properties like non-immunogenic, biocompatibility, good biodegradability it is also an excellent molecular targeting moiety as it readily recognizes CD44 receptors on tumors. **^[28]^**The chemically active molecules on HA could provide different modification strategies. Furthermore, this polymer could be immobilized onto light responsive GNRs using different methods such as electrostatic interaction, one-pot ligand exchange, and layer-by-layer. **^[29, 30]^** Upon NIR irradiation the GNRs induces hyperthermia release of encapsulated anticancer drug by breaking covalent bond between polymer and anticancer drug. **^[31, 32]^** Therefore, designing ideal nanoplatform to achieve excellent biocompatibility is the key role to introduce robust delivery system so as to minimize undesired effect. Herein, we prepared a multifunctional platform (MTX-HA@GNRs) by integrating different components namely anticancer drug methotrexate (MTX), GNRs as photoresponsive agent, drug carrier and targeting-specific moiety HA. The MTX is an antimetabolite genotoxic component which essentially inhibits mechanism of DNA, RNA and slows the growth of proliferating cancer cells. **^[33, 36]^** Whereas, natural polymer HA have been researched throughout the decades because of their easy adaptation in biological system. **^[35, 34]^**Besides biological properties like non-immunogenic, biocompatibility and molecular targeting moiety as it actively recognizes CD44 receptors on tumors. **^[36]^** The chemically active repeating units of hydroxyl and carboxyl molecules on HA allowed covalent conjugation with MTX via “Ester coupling” reactions. This prodrug conjugate supported immobilization onto GNRs using electrostatic interaction. **^[37]^** Initially, GNRs were modified with cationic ligand to enhance water dispersion simultaneously surface charge of TMA adapted electrostatic interaction with anionic MTX-HA. **^[38]^**The preliminary heat from plasmonic GNRs upon receiving NIR produces localized heat and accelerated hydrolytic cleavage of the ester bond between MTX-HA. **^[39]^** This CHT-PTT channeled together demonstrated MTX release from TGNRs@MTX-HA. Additionally, the affinity of HA towards CD44 receptor expressed on tumors guided localization of MTX. **^[40]^** As a result, this designed nano-platform provides several advantages like **1)** Safe and prolonged circulation **2)** Dual targeting i.e. active targeting and biocompatibility due to HA and passive targeting from GNRs; **3)** Finally, the controlled release of drug MTX was monitored via hydrolytic cleavage of ester linkage in acidic tumor environment due to elevated heat from NIR.**^[41]^** This research marks a new strategy for potential delivery of all types of hydrophobic and hydrophilic drugs also provides one of the promising platforms that constrain efficient combination therapy comprised of CHT and PTT.

## 2. Materials and analytical methods

### 2.1. Materials and Equipment

#### a) Materials

Chloroauric acid tetra hydrate (HAuCl_4_.4H_2_O ≥99.9%), hexadecytrlimethylammonium bromide (CTAB ≥ 96%), sodium borohydride (NaBH_4_, 99%), silver nitrate (AgNO_3_ ≥ 99.9%), L-ascorbic acid (AA ≥ 99.0%), 11-bromo-1-undecanol (98%), triphenylmethanethiol (97%), methanesulfonyl chloride (MsCl; 98%), trifluoroacetic acid(≥99%), triisopropylsilane (98%), trimethylamine (≥ 99.5%), and triethylamine (≥ 99 %) were purchased from Sigma Aldrich. For further synthesis, Rhodamine B used as fluorescent probe, Methotrexate (MTX, ≥ 98 %), Hyaluronic acid (HA, MW 10KDa), Dialysis membrane (2KDa), 4- Dimethlaminopyridine (DMAP, ≥ 99 %), N, N’-Dicyclohexylcarbodiimide (DCC, 99%, Dimethyl sulfoxide, Formamide, for synthesis were purchased from Sigma-Aldrich and Tokyo Chemical Industry (TCI), Thermo Scientific respectively. To conduct release experiment Phosphate buffered saline solution (PBS) was purchased from Sigma-Aldrich. Ultrapure (D.I) was used for conducting release experiments. All cell reagents for *invitro* studies, Phosphate buffered saline solution (PBS), Dulbecco modified Eagle’s medium (DMEM), fetal bovine serum (FBS), pen-strep were purchased from Sigma Aldrich. The cell viability was quantified using 3-(4, 5-dimethylthiazol-2- yl)-2, 5-diphenyltetrazolium bromide (MTT) which was purchased from Sigma Aldrich.

#### b) Equipment

To conduct NIR release experiment infrared camera (Fluke, USA) was used. H^1^NMR was used to analyze the chemical synthesis ECX-400 instrument (JEOL, Japan). To characterize, the TEM image was obtained via JEOL 1000CX (JEOL, Japan) operating at 80keV and the modified GNRs were monitored using a Nicolet iS 10 FT-IR spectrometer (Thermo Scientific, USA). The charge density was acquired using Zeta potential analysis by Dynamic Light Scattering (DLS) Zetasizer. The spectral absorption was quantified by Neosyss-2000 UV-vis spectrophotometer (SCINCO, USA) and Fluorescent spectrometer QM-400 (Horiba Scientific, USA). The cell counting and viability was measured via microplate reader (Tecan infinite series 200, Germany). The cell sorting after staining was obtained by (fluorescence-activated cell sorting (FACs) caliber, BD Bioscience).

### 2.2. Preparation of CTAB-Gold nanorods (CGNRs)

CTAB-GNRs with peak absorption 780nm were prepared according to the seed-mediated growth protocol by using familiar surfactant CTAB. Briefly, a seed solution was made by adding 7.5ml of CTAB (0.1M), 250µl of HAuCl_4_ (0.01M), and 800 µl ice-cold of (0.01M) NaBH_4_. The solution was stirred for 2-3 h to get a brown yellow solution. The GNRs growth solution of total volume 30ml which contained 1.8ml of 0.01MHAuCl_4_, 180 µl of 0.01M AgNO_3_, 288 µl of 0.1M AA were gently prepared and then 130 µl seed solution was introduced, the solution was kept without disturbing at 35 ^0^C for 24h.The excess of CTAB from GNRs solution was removed using centrifugation at 13000 rpm and stored for further experiments.

### 2.3. Preparation of monothiol TMA-GNRs (TGNRs)

CTAB-GNRs were re-centrifuged at least twice at 13000rpm for 20 min to remove the excess of CTAB before use. Synthesized TMA was covalently attached to the GNRs surface via thiol affinity bonding following the one-pot ligand place-exchange method. The whole process was applied in D.I water due to good solubility of TMA. To prepare the exchange of TMA-GNRs, Solutions of TMA ligand (0.25ml, 10mm) were added drop wise in GNRs solution (7.5ml) followed by gentle stirring for 72 h. The resulting complex was centrifuged at 13000rpm for 20 min to remove the excess of TMA ligand and the final concentration was fixed by Beer- lambert law and molar extinction coefficient.

### 2.4. Preparation of TGNRs@MTX-HA

RHO.B-HA is coated onto TGNRs using electrostatic interaction. Briefly, Aqueous suspension of TGNRs (20.3×10^-9^M, 3.5 mL) in D.I was introduced drop wise into RHO.B-HA aqueous suspension (15mg, 10ml) resulting mixture was covered and allowed for stirring for 48h, this whole procedure was applied in dark condition. RHO.B-HA@TGNRs was collected and dialyzed against D.I to remove free RHO.B, Monomer or dimer of HA, applied for lyophilization. To check stability, it was redispersed in D.I and peak absorption with LSPR and TSPR shift was confirmed by UV-vis spectrophotometer.

### 2.5. Evaluation of thermal elevation efficiency of TGNRs@RHO.B-HA

To elucidate the concentration effect of suspension on NIR triggered PTT conversion ability and elevation of Temperature, different concentration ranging (0.2, 0.4, 0.6, 0.8, 1.6 nM) and different laser power ranging (0.2, 0.4, 0.6, 0.8, and 1.6 W/cm^2^) was prepared and subjected for testing NIR. Briefly, suspension of TGNRs@RHO.B-HA with different concentrations was added to 0.5ml PBS in a mini dialysis tube and was positioned in the center of the NIR laser beam. The power was set at 1.6 W/cm^2^, it was irradiated for 15min using NIR laser. Thermal elevation from laser irradiation monitored using an infrared camera for every 1 min set time point. Temperature changes and photostability of this conjugate were clarified by repeating 3 times to minimize possible errors and results were noted.

### 2.6. Thermal responsive release of model drug RHO.B from TGNRs@RHO.B-HA

To evaluate temperature stimuli release of RHO.B at different irradiation times, TGNRs@RHO.B HA was dispersed in PBS, later 0.5ml of a solution of TGNRs@RHO.B-HA was added to an Eppendorf tube containing 0.5ml PBS. The tube was placed in the center to acquire an accurate beam from a laser, the power 1.6 W/cm^2^was irradiated by NIR laser (808nm) for 30min at 5 intervals. After irradiation, the samples were centrifuged at 13000 rpm for 15 min and the amount released free RHO.B was measured via Fluorescence spectroscopy.

### 2.7. Evaluation of Activation Energy to study Hydrolysis of Ester Bonds

To calculate the kinetic release and activation energy of the sample, the solution of TGNRs@RHO.B-HA was prepared in PBS and distributed in a dialysis tube. The sample tube was then placed at different temperatures (25, 37.5, 50, and 70 °C) which were maintained in a water bath. At beginning consecutively for 3 h, 0.5ml sample was withdrawn and replaced with 0.5ml of fresh PBS and it was continued until a saturation point was observed. The collected sample was quantified via spectrofluorometer and activation energy of the hydrolysis of ester bond was calibrated and the graph was plotted.

### 2.8. *Invitro* experiments

#### 2.8.1 Cell culture

Cell lines HeLa cells and HEK293 cells were obtained from ATCC the optimal parameter was maintained for culturing cells. Initially DMEM medium was prepared by adding 1% penicillin and streptomycin later supplemented with 10% fetal bovine serum (FBS). After successful addition of DMEM media culture flask was incubated at 37^0^C in 5% CO_2_ humidified incubator and cell lines were used for further assays after confirming cell density.

#### 2.8.2 Cell uptake and Cytotoxicity studies

To investigate active cellular uptake different concentrations of TGNRs@MTX-HA, TGNRs@HA and TGNRs were incubated HeLa and HEK293 cells and analyzed by laser confocal fluorescence microscopy (LCFM). Briefly, cells were seeded with 1000 cells/well density in the LCFM 24 well culture plate and incubated at 37^0^C, 5% CO_2_ incubator to achieve 80% confluence of cells. The cells were treated with 50μM and 100μM of TGNRs@MTX-HA, TGNRs@HA, and TGNRs respectively and incubated for 3h, 6h, 12h and 24h and irradiated with laser power of 1.6 W/cm^2^ except blank. To remove cell debris washing step was followed by Dulbecco phosphate buffered saline (DPBS pH 7.0) for 3 times, LCFM was performed and fluorescence images were obtained using above method.

To determine cytotoxicity and cell viability the HeLa and HEK293 cells were seeded with density of 1000cells/well in 96well plate. Different concentration of TGNRs@MTX-HA, TGNRs@HA and TGNRs (0.0625, 0.125, 0.25, 0.5, 1, 2.5, 5nM) in D.I was added and allowed for 24h of incubation. The cells were irradiated with laser power 0.6 Wcm^-2^ for 5 min, except media control well. The MTT reagent was added into the cells and incubated for 3h, afterwards the formed formazon crystal was diluted using DMSO (100μl in each well) and absorbance was obtained using microplate reading at 570-590nm.Cell viability was calculated using below formula: **(mean O.D of treated wells-mean O.D of control wells)/mean O.D of untreated wells-mean O.D of medium control well)*100**

## 3. Results and discussion

Nanomaterial-assisted PTT has potentiated treatment method in the oncology field. Thus the importance of inventing Gold based nano-platform using active targeting moiety has marked functional recognition which not only improves cancer-targeting but also provides controlled t release of anticancer drug. **^[42]^** For example, Yang’s research group prepared camptothecin coated GNRs for CHT-PTT, the results showed that micro fabricated GNRs not only accumulated passively in cancer cell but also allowed drug release responding to NIR. **[43]** Such nano-platforms that use light responsive nanomaterial as GNRs deeply facilitates thermal ablation of cancer cells by passive targeting effect, whereas negligible effect to the surrounding healthy cells. **^[44]^**

Scheme 1 shows our approach to prepare GNRs based nanocarrier TGNRs@RHO.B-HA briefly, owing to high quenching fluorescence property we incorporated model drug Rhodamine B using covalent bond to hyaluronic acid via hydrolytic ester bond, and further this conjugate favored electrostatic interaction with GNRs.This electrostatic interaction is the main driving force which was possible due to surface modification of CTAB-GNRs with thiol ligand N, N, N-trimethyl- 11-(tritylthio) undecan-1-aminium (TMA). In addition the known thiol (-SH) ligand was initially synthesized using organic synthetic method. TMA enhances GNRs dispersion in water and positive charge promoted electrostatic interaction. **^[45, 46]^**

**Figure 1.**
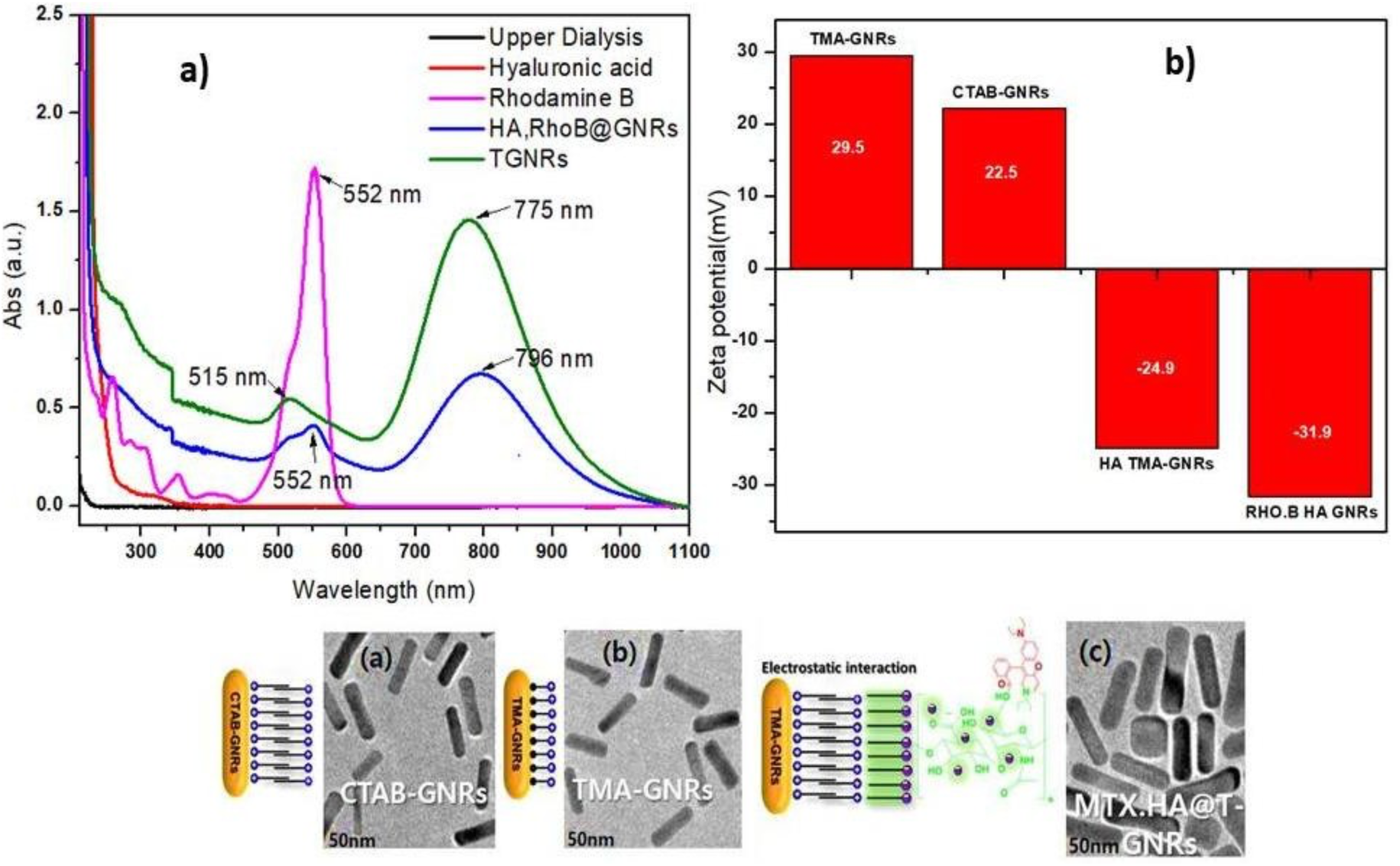
**(a)** UV-vis spectra of CTAB-GNRs, TMA-GNRs (TGNRs), HA@TGNRs, HA- Rho.B@TGNR **(b)** Zeta potential of CTAB-GNRs, TMA-GNRs (TGNRs), HA@TGNRs, HA- Rho.B@TGNRs and Figure 1. TEM image **a)** CTAB-GNRs, **b)** TMA-GNRs (TGNRs), **c)** HA@TGNRs, d) HA-Rho.B@TGNRs.

To address photothermal properties in regard to NIR range tunable longitudinal SPR (730-800 nm) fabrication of CTAB-GNRs was achieved by classic seed-mediated method; In general, this method proceeds for 2 h of seed growth followed by 10 h of rod formation. **^[47]^**Higher yield is obtained of GNRs with controlled Plasmon resonance and aspect ratio by adjusting the ratios of surfactant CTAB which also prevents aggregation of particles. The excess of CTAB attached on the surface of GNRs was eliminated by repeated centrifugation. Although, this method is used frequently due to its promising yields but it fails to maintain stability of GNRs in solvent system. In order to provide aqueous stability, GNRs were modified with TMA ligand using “one-pot ligand exchange method “in deionized water (D.I).

In this case” Thiol (-SH)” ending cationic ligand TMA was pre-synthesized as gold-based nanoparticle shows strong affinity towards “sulfur” to form“GNRs-Sulfur”. The replacement of toxic surfactant CTAB with TMA facilitated conjugation to anticancer drug MTX-HA using physical adsorption, illustrated in **Figure 1**. The constructed nanoplatform after successful ligand exchange was analyzed using Uv-vis spectroscopy and TEM, results of TEM have confirmed the size ∼60nm and aspect ratio 3.6±1.7 respectively.The increase in size of GNRs after modification with MTX.HA is possibly due to hyaluronic acid.Whereas, Uv spectrum results demonstrated the shift of TMA-GNRs strong longitudinal plasmon (LSPR) band peak from 745-800nm, compared to CTAB-GNRs which was 750nm. This conformational information was the mainstay to further investigate plasmonic activation using NIR. **Figure 1(a) and (b)**

For potential nano-carrier the possible mechanism that exhibits endocytosis of nano-agent depends on the charge, size, and biocompatibility. Basically, GNRs are fabricated using bilayer CTAB during its formation however such surfactants are cytotoxic for medical application. To enhance bioactivity, GNRs are tuned with biocompatible active molecules namely polymers, receptors etc. In present study, the anionic glycosaminoglycan HA the known polymer owing to high bioavailability is immobilized onto cationic TMA-GNRs using electrostatic interaction by simple round-trip ligand coating method. The physical adsorption of MTX-HA on GNRs via electrostatic attraction broadens the spectrum peak at 500nm and 800nm.

The corresponding electrical charge of GNRs increased by two-fold. The zeta-potential analysis of different complexes such as CTAB-GNRs, TMA-GNRs, HA-GNRs, HA-MTX@GNRs showed variation in surface charges as well as a gradual shift in size. The presence of CTAB appeared ± 22.2mV, after covalent conjugation of TMA the values increased + 39mV whereas, reversal of charge was found after coating with HA and HA-MTX because of anions -24.9mV and -41.6mV respectively. Interestingly, this surface modification provided the evidence about maximum removal of toxic CTAB and enables to construct promising biomaterial. **Figure 1(c)** The functionalization of surface chemistry of GNRs is very important finding for various biomedical applications. The strong electrostatic interaction between the TMA and HA-MTX or RHO-HA provided the better option to prepare conjugate which was analyzed by FTIR spectra. Initially, different samples were prepared i.e. CTAB-GNRs, TMA-GNRs, and TGNRs@RHO- HA. The FT-IR spectra of respective samples are shown in Figure S. The peaks of C-N stretching of CTAB from CTAB-GNRs appeared in the range 2500-3000 and 1400-1490cm^-1^. After covalent attachment of TMA ligand onto GNRs, the presence of additional C-N stretching spectral band was observed around 1630-1690cm^-1^ this mean difference indicated the replacement of cetrimonium bromide CTAB by TMA. The existence of C=O spectral stretch around 1580-1750 cm^-1^ expectedly occurred from ester bond between MTX-HA. This result of FT-IR endured the successful functionalization of GNRs. **Figure S2**

To confirm clinical application using external stimuli, especially NIR irradiation is a potent to release the drug in sustained manner in our system. To endow the thermal-dependent release we prepared analogue using fluorescence probe Rhodamine (RHO.B) dye as model drug and conjugated to HA using esterification coupling reaction. Further this conjugate was electrostatically adsorbed onto TMA-GNRs. As RHO-B shows strong absorbance intensity than MTX thus free traces of RHO.B was easily quenched using fluorescence spectrometer even with low concentration. **(Figure 3a)**

The free traces of TGNRs@RHO.B-HA were recorded with different concentrations and laser powers with respect to time. For instance, varied concentration 0.2, 0.4, 0.8, 1.6 nM of TGNRs@RHO.B-HA was irradiated with laser diode 808nm, power 1.6 W cm^-2^ for 15 min and temperature was noted using infrared camera for every 1min, as a control, there was no significant increase in temperature in phosphate buffered solution (PBS) without addition of GNRs. **Figure 2(c)** But competitive increase of temperature from 35 °C to 65 °C was obtained from 0.2, 0.4, 0.8, 1.6 nM of TGNRs@RHO.B-HA. The fluorescent traces of free RHO-B were recorded using fluorescence spectrometer. **Figure 3(b)**

**Figure 2.**
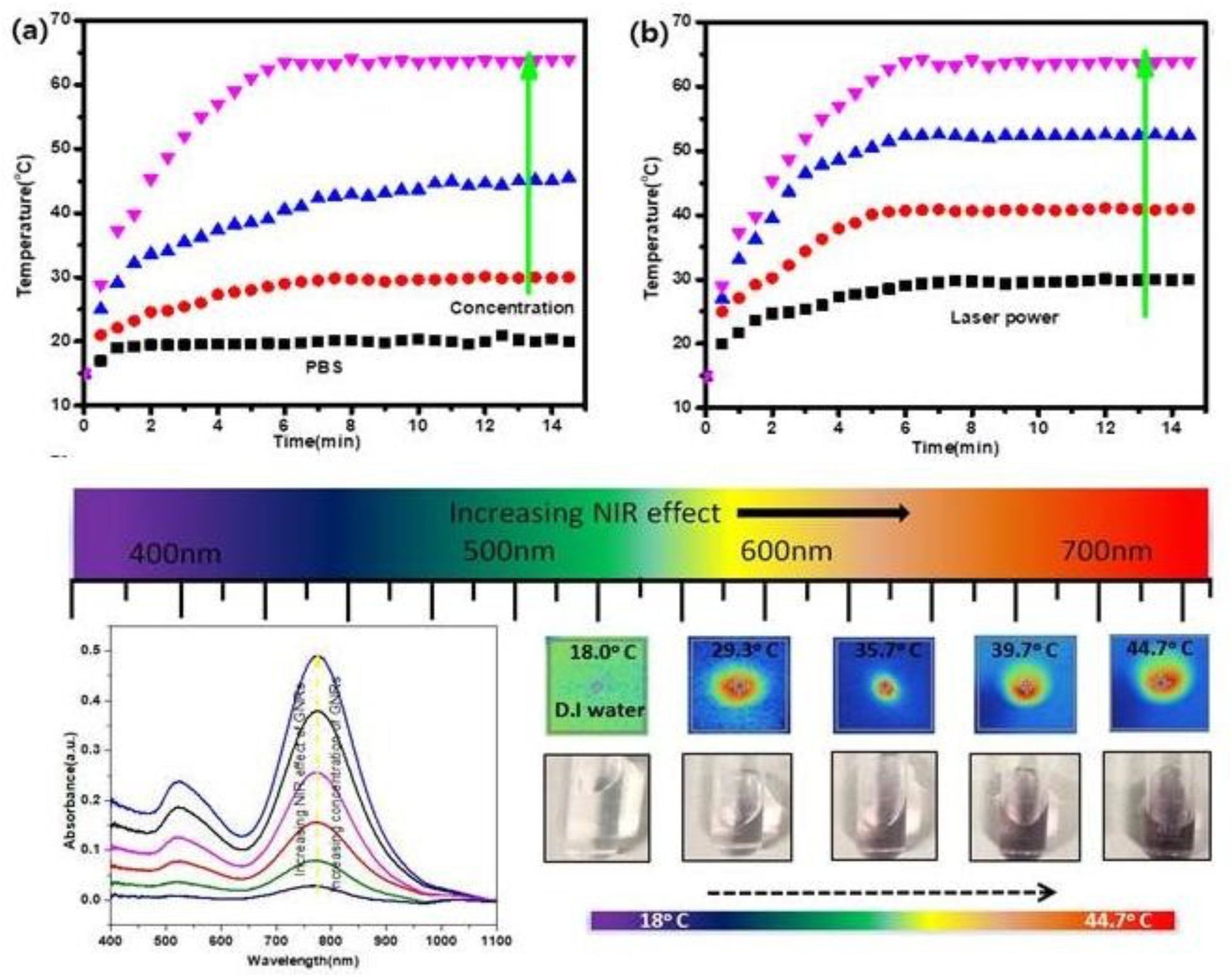
Analysis of temperature elevation **(a)** depending on concentration (0.15, 0.5, 1, 2.5 nM) with laser power 1.6W/cm^-2^ for 15min **(b)** depending on laser power (0.8, 1.1, 1.6W/cm^-2^)

In addition to specify controlled release of our nanoplatform, TGNRs@RHO.B-HA solution of 0.8nM was irradiated with different laser power 0.4, 0.6, 0.8 and 1.6 W cm^-2^. The results effectively the increase in temperature which determined the effect of concentration of nanomaterial GNRs is an outstanding source of thermal increase associated with laser power density. Thus, this study promoted efficacy of programmed drug release pattern from our nanoplatform TGNRs@RHO.B- HA. **Figure 2(b)**

The controlled release of drug payload using rate constant from the nanoplatform enhances the deeper understanding of programmed release of encapsulated drug from the system. The chemical bond ester is microenvironment pH sensitive, to study release rate of RHO-B from TGNRs@RHO.B-HA by hydrolysis of ester bond was carried using concentration 5mg/1.5ml in PBS added to dialysis bag. The temperature was maintained 25, 37, 50, and 70°C and dialysis bag placed in each water bath. For 24h, the 500 μl was collected and replaced with fresh PBS. This result provided the hydrolysis rate of ester bond with respect to time and temperature by first order kinetic which is concentration dependent indicating sustained release of model drug could be controlled **Figure 3(a)**. Further the activation energy (Ea) was calculated as 36.51KJ mol^-1^, which is based on hydrolysis rate of ester bond. The linear release relation of free RHO.B provided additional information of mechanism to describe the amount of energy used to dissociate the ester bond over the increasing time. **Figure 3(a) and (b)**.

**Figure 3. a).**
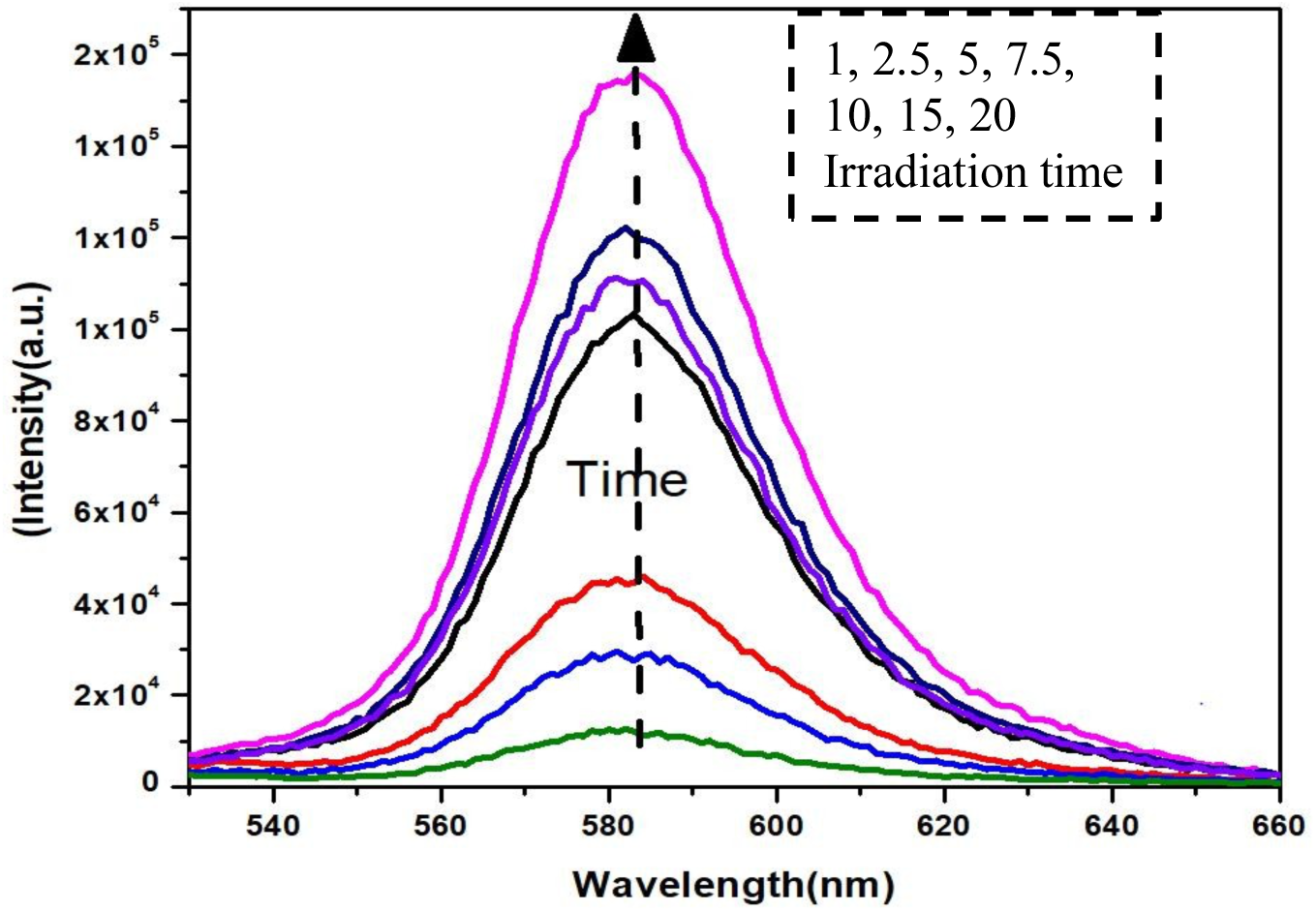
Accumulative concentration of released free RHO.B from HA-RHO.B@TGNRs at variable time with laser irradiation at 1.6W) in PBS

**Figure 3. b).**
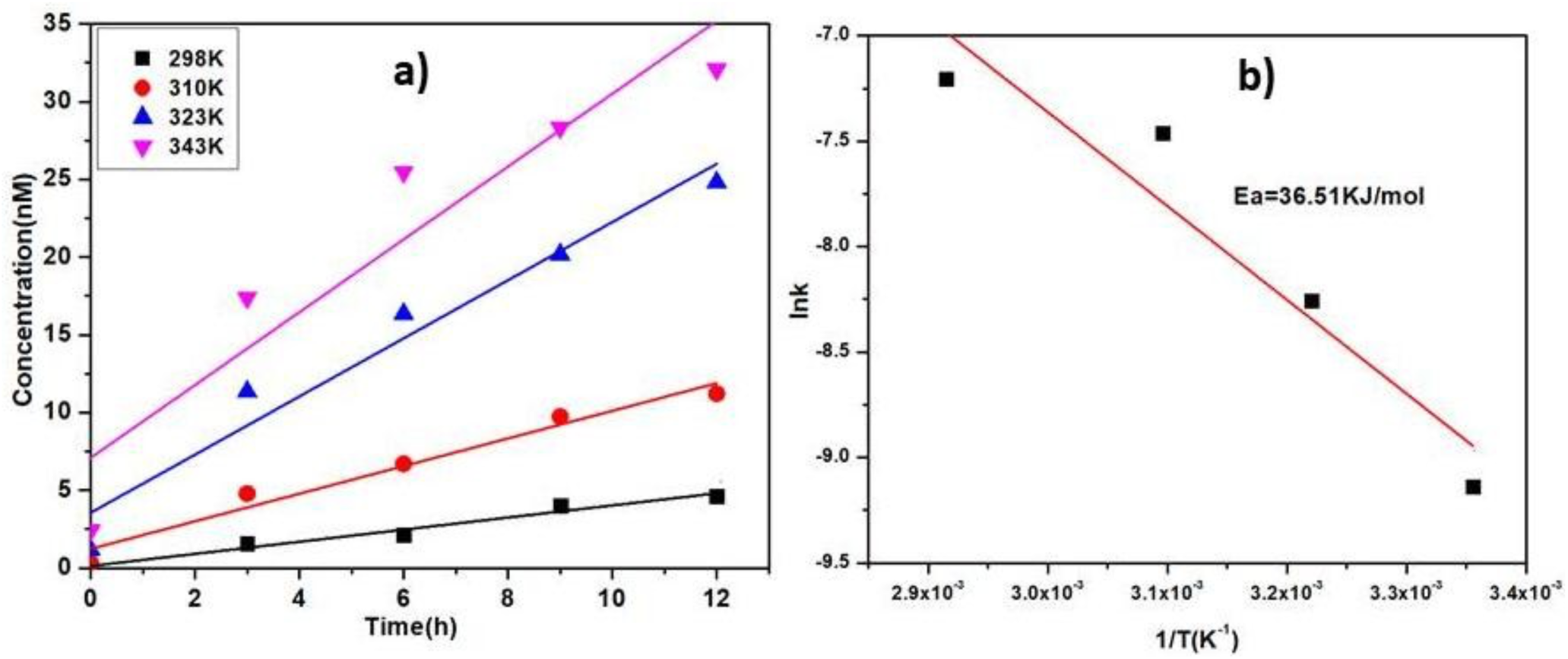
Accumulative concentration of released free RHO.B from HA-RHO.B@TGNRs at variable temperatures (298, 310, 323, and 343 K) and **b)** Activation energy of hydrolysis rates of model drug RHO.B from HA-RHO.B@TGNRs in PBS.

active targeting ability of HA towards CD44 receptor suggested cell-specific uptake when added with GNRs and NIR respectively. To elucidate the active cellular uptake, HeLa cells where preincubated with free HA to block the CD44 receptor for 2h, after that different concentration of TGNRs@RHO.B-HA was added and incubated at different time. In pretreated HA group there was no mean flourosence found may due to blocking of CD44, whereas as non-treated group the displayed strong fluorescence emission and excitation of free RHO.B from TGNRs@RHO.B-HA however non-coated GNRs found to be escaped from cellular uptake, this indicated the active targeting ability of HA towards CD44 receptor over expressed in HeLa cells **(Figure 4a)**

**Figure 4. a).**
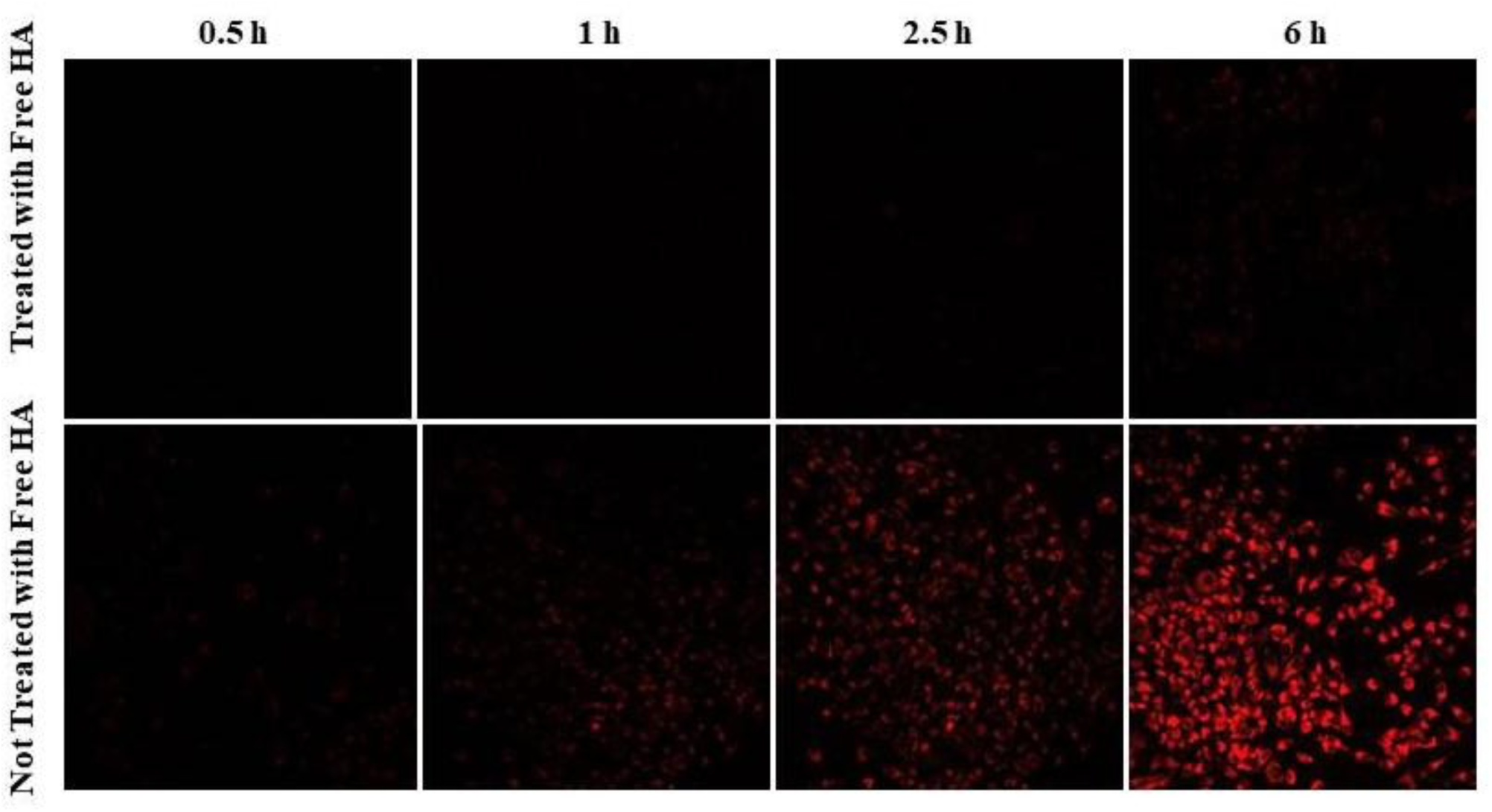
Uptake difference for receptor specific analysis before (-) and after (+) addition of free HA for HA-Rho.B.

The localized thermal activation of GNRs passivized nanoplatform the accumulation of nanocomplex stimulated by EPR effect in the tumor cells attributes certain drawbacks like low uptake, carcinogenic to healthy cells and irreversible drug waste. The unmet result from our previous research expectedly limits the use of GNRs without the further modification with known biocompatible moiety. The GNRs showed passive internalization of anticancer drug mediated by hyperthermia when irradiated with different laser powers but excellent plasmonic activation was observed with 1.6 W cm^-2^laser powers. The current nanocomplex comprised of biopolymer HA with abundant functional group for further binding facilitated formation of TGNRs@MTX-HA and model anticancer TGNRs@RHO.B-HA. In presence of GNRs (-) the uptake of RHO.B-HA was enhanced with external stimuli i.e. 1.6 W cm^-2^ laser power. Whereas in absence of GNRs (-), the uptake was reduced may be due to receptor specific endocytosis which is time dependent. Overall the **(figure 4.b)** states the combination effect is more promising over single therapy.

**Figure 4. b).**
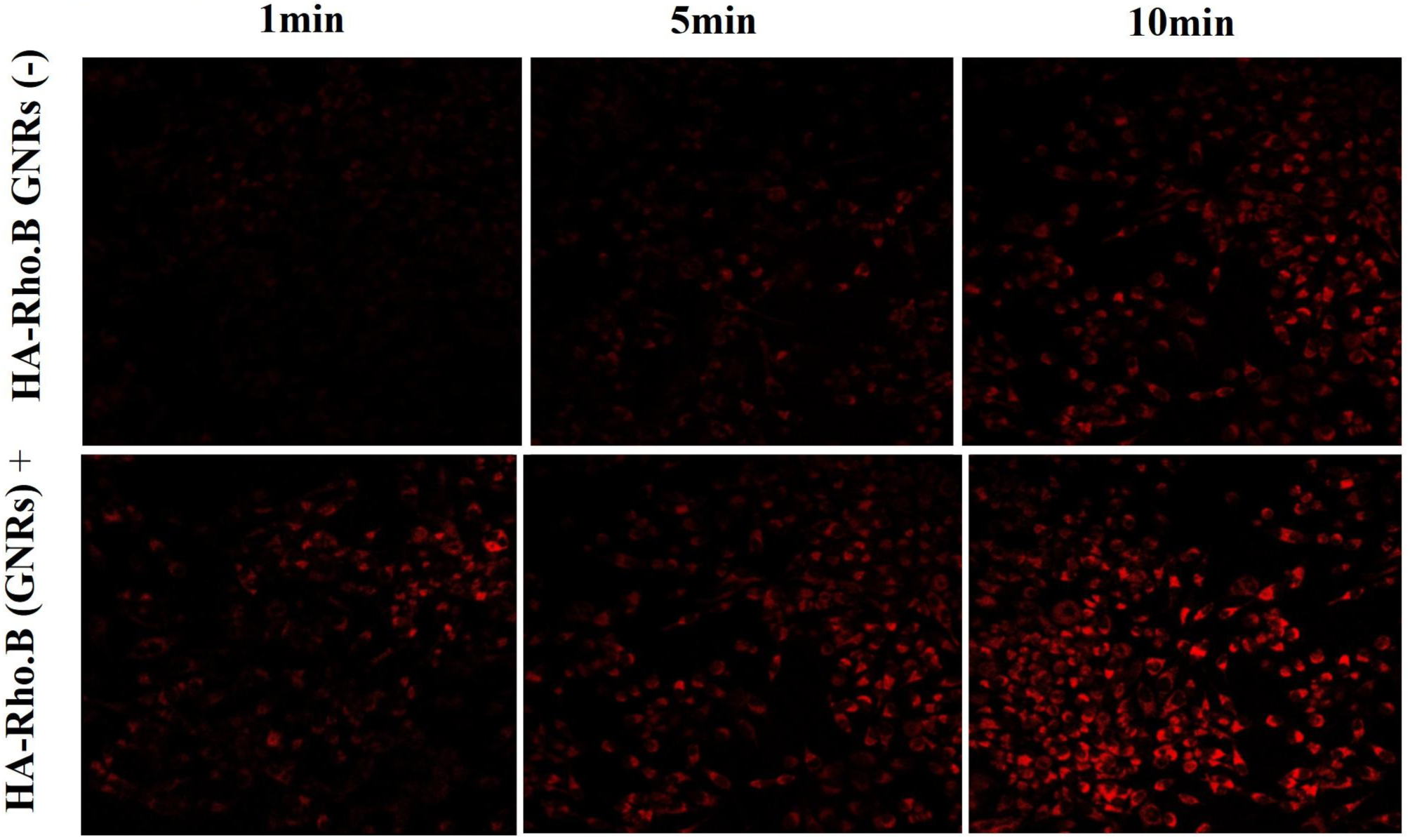
Uptake difference depending on laser irradiation at 1.6W for 1, 5, 10min for in absence and presence of GNRs for HA-Rho.B.

The cellular uptake of TGNRs@MTX-HA could be enhanced by increasing laser power and exposure time **(Figure 5a,c)**. Designing bio-absorbable nanocomplex structured with biopolymer favors non-immunogenic reaction for normal cells in cancer environment. **[48]** Specific receptors help to target malignant cells and induce non-toxic effect for the surrounding normal cells. In this work we assessed the biocompatibility of TGNRs@MTX-HA, TGNRs@HA and TGNRs. Briefly, The results obtained significantly hold low toxic effect with cell viability 97% NIR (+) for 0.05nM and 62% NIR (+) for 5nM on NIH3T3 cell. Whereas, for TGNRs@MTX-HA, 0.05nM showed 88% cell viability and 46%, 5nM with NIR irradiation when compared to control from graph. This experiment helped to demonstrate the cell death occurred by endocytosis of GNRs and drug accumulation after thermal cleavage of ester bond between HA-model drug. The results for biocompatibility of the TGNRs@HA expectedly showed low cytotoxicity when incubated without anticancer MTX drug in NIH3T3 cell. For combined CHT-PTT and cytotoxicity effect the MTT assay was performed in presence of NIR by incubating with HeLa and NIH3T3 cells with variable concentration of TGNRs@MTX-HA, TGNRs@HA, and TGNRs. **[49]** Upon irradiation with NIR laser diode for 5, 10, 15 min at power 0.8W cm ^-2^ there was no significant morphological change after observing by LCFM. The EC50 value was calculated and estimated that the minimal concentration required inhibiting the growth **(Figure 5d)** the comparative studies between (-) CD44 U87 and (+) CD44 Hela cells showed decrease in cell viability of HeLa cells upto 95% depending on combine CHT-PTT with laser power 1.6W and time 1, 10, min irradiation. We also visualized the increased cytotoxic effect of TGNRs@MTX-HA, as compared to TGNRs@HA and TGNRs in HeLa cells.

**Figure 5.**
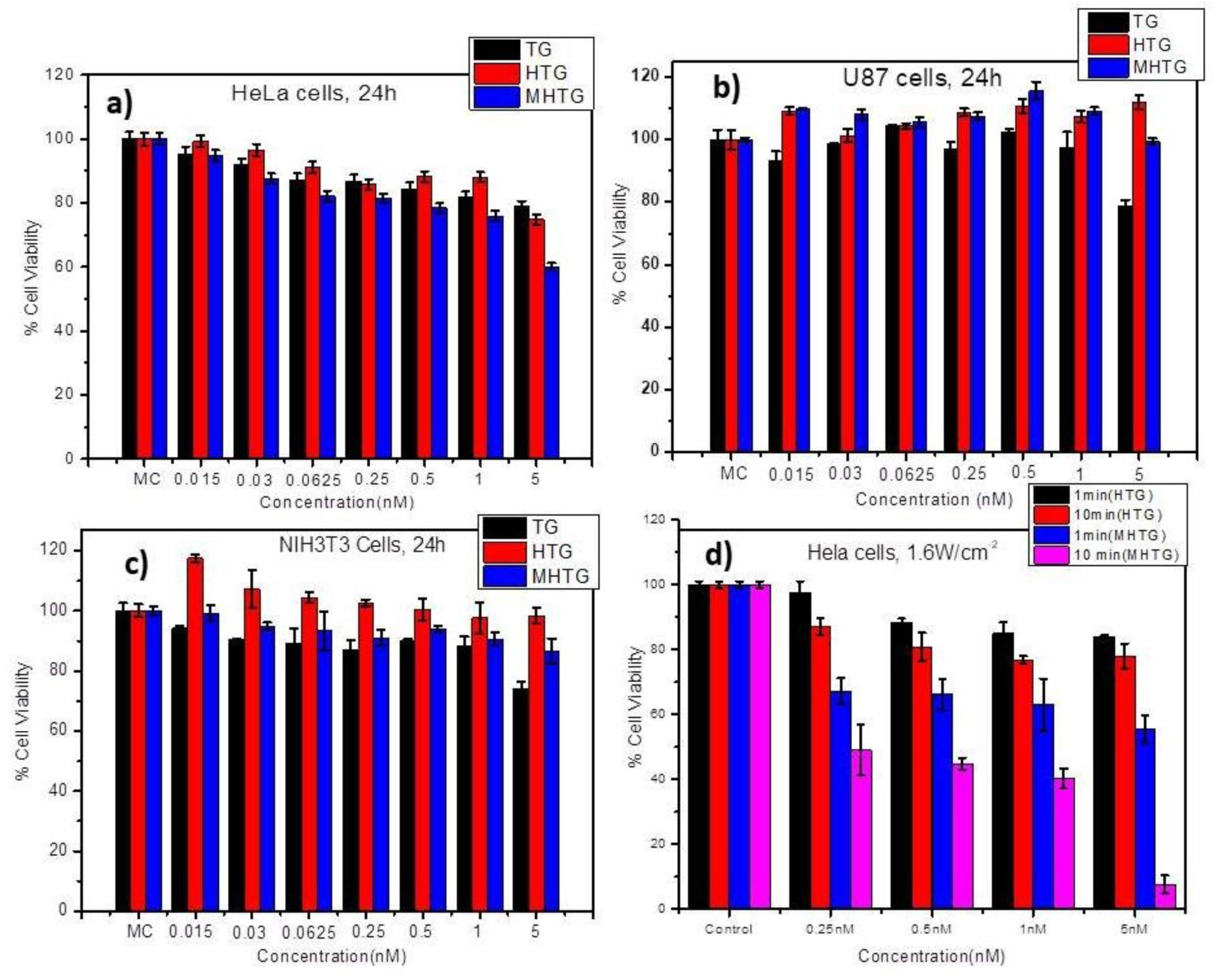
*Invitro* studies **a)** and **b)** without NIR using TG, HTG, MHTG for receptor specific cell toxicity. **c)** Biocompatibility of TG, HTG, MHTG in NIH3T3 without NIR and **d)** CHT-PTT using 1.6W NIR for 1, 10min using TG, HTG, MHTG.

## 4. Conclusion and Future work

GNRs in multi-disciplinary field especially for cancers have demonstrated complementary to the traditional methods in regard to optically active therapy. The multivariate properties such as surface modification of GNRs with synthetic ligand help to increase bio-distribution and stability in the solvent system. On addition of HA biopolymer promisingly supported the active targeting effect and localization of hydrophilic/hydrophobic anticancer drugs.

However development of nanoplatform with aim to provide high loading efficiency, good biocompatibility with easy elimination of nanoagents from bodily system is still ongoing process. Meanwhile, the experimental results of our designed robust nanoplatform have improved previous limitations. As MTX possess low quenching intensity so we synthesized RHO.B-HA to check release profile of free RHO.B using NIR light irradiation upon hydrolysis of ester. The DCC and DMAP coupling facilitated the esterification between free carboxyl group of MTX and hydroxyl of HA and analogue RHOB-HA.Using simple ligand exchange method we successfully replaced the surface chemistry of GNRs.Furthermore using physical force such as electrostatic interaction exhibited coating of MTX-HA and model drug RHO.B-HA onto surface of GNRs. The heat sensitive covalent ester bond in cancer microenvironment allowed novel insights to perform drug release on responding to localized hyperthermia generated from NIR responsive GNRs.Whereas HA provided discrimination between cancer and normal cell with active cellular uptake due to affinity towards CD44 receptor in cancer environment. Interestingly it was found that nanoformulation MTX-HA@TGNRs excellently encapsulated drug, ensured biosecurity because of targeting effect and lastly provided localized thermal cytotoxicity to cancer. This new strategy could be biological target to link anticancer drug associated with synergistic effect when combined CHT and PTT for various medical applications in futu

## Supporting information(S)

## (S1) a) Synthesis of Rhodamine B (RHO.B) -Hyaluronic acid (HA)

Rhodamine-B was conjugated chemically to hyaluronic acid (HA) by the ester bond formed via DCC and DMAP coupling **(S1)**. In brief, HA (150mg, 0.015mmol) was dissolved in formamide (2mg/ml) and allowed to stir for complete mixing and it was diluted with 7ml of dimethyl sulfoxide (DMSO). Separately Rhodamine-B (94mg, 0.205mmol), DCC (175mg, 0.848mmol), DMAP (115mg, 0.9417mmol) were dissolved in 10ml of DMSO, followed vigorous stirring for 45min- 1hr for the activation of carboxyl group of Rhodamine-B. Resulting activated complex was added drop wise to the HA solution and reaction was allowed in dark for 24 h at room temperature. The obtained crude product was centrifuged at 5000 rpm for 15min to remove precipitated dicyclohexylurea residue. The supernatant was proceeded for dialysis against deionized water to remove unattached Rhodamine-B using a membrane (MW cutoff=1000 Da).The complex was lyophilized and stored for further experiments. The physico-chemical analysis was obtained using ^1^H NMR solvent D_2_O δ

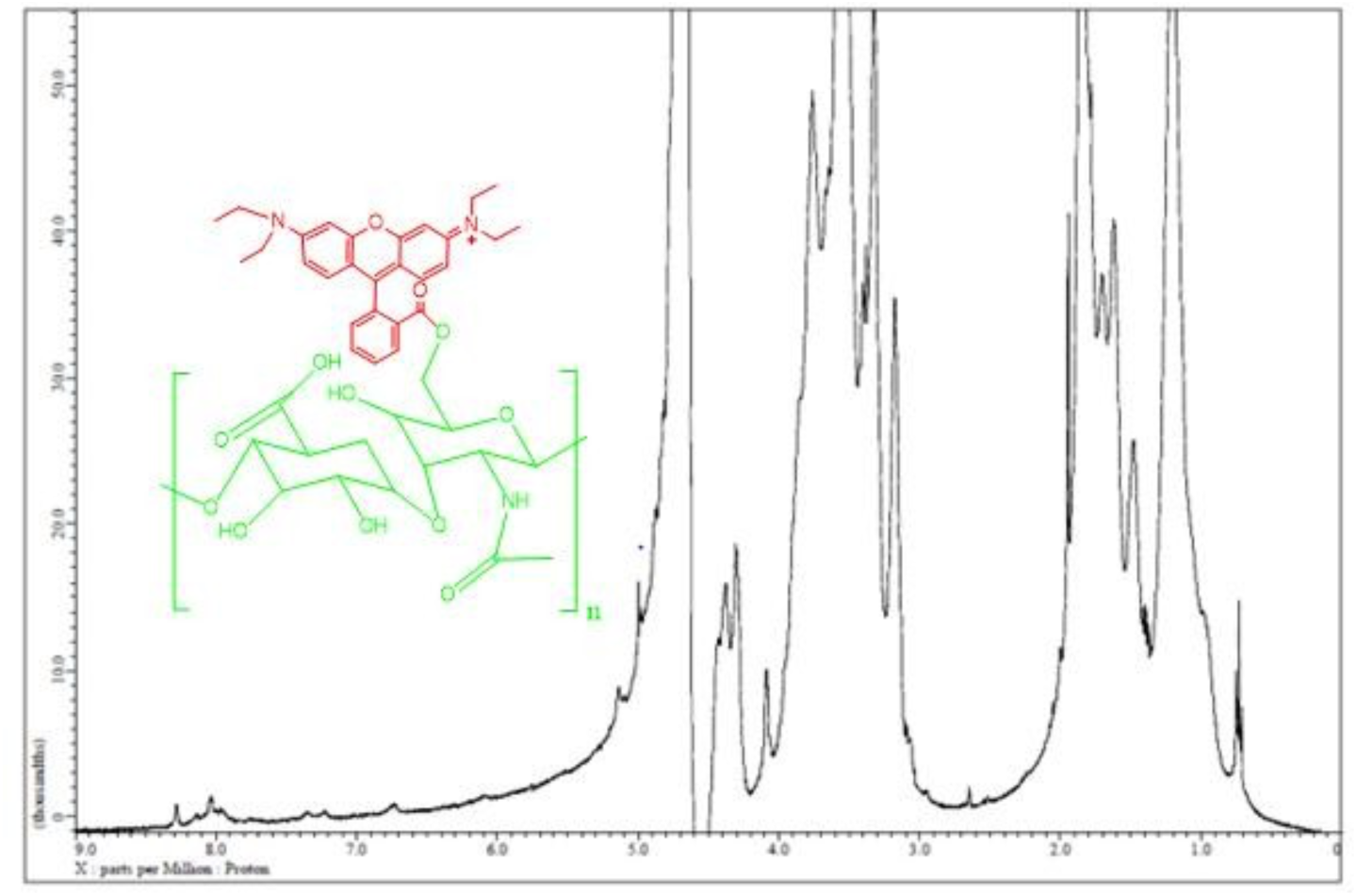

## b) Synthesis of Methotrexate (MTX)-Hyaluronic acid (HA)

MTX was conjugated chemically to hyaluronic acid (HA) by the ester bond formed via DCC and DMAP coupling **(S1)**. In brief, HA (150mg, 0.015mmol) was dissolved in 35ml of formamide and allowed to stir for complete mixing and it was diluted with 7ml of dimethyl sulfoxide (DMSO). Separately MTX (94mg, 0.205mmol), DCC (175mg, 0.848mmol), DMAP(115mg, 0.9417mmol) were dissolved in 10ml of DMSO, followed vigorous stirring for 45min-1hr for the activation of carboxyl group of MTX. Resulting activated complex was added drop wise to the HA solution and reaction was allowed in dark for 24 h at room temperature. The obtained crude product was centrifuged at 5000 rpm for 15min to remove precipitated dicyclohexylurea residue. The supernatant was proceeded for dialysis against deionized water to remove unattached Rhodamine-B using a membrane (MW cutoff=1000Da). The complex obtained was lyophilized and stored for further experiments and analysis. The chemical shift in ^1^H NMR δ ppm shift for MTX-HA in D_2_O observed as (2.836, 7.096, 7.115, 7.129, 7.916, 7.938, 7.978, and 8.851) compared.

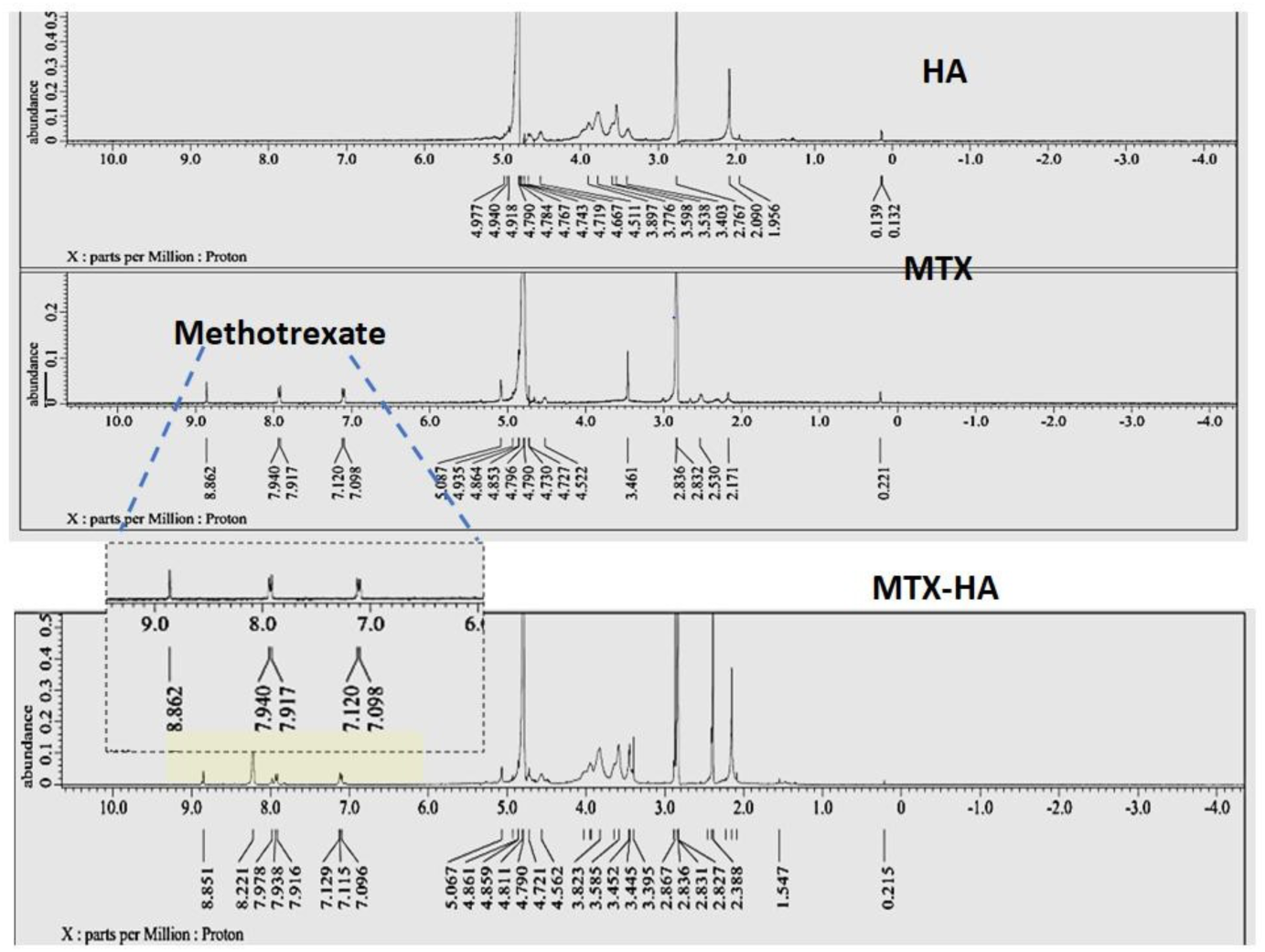

## (S2) Uv-vis and DLS of (HA, MTX, HAMTX, HAMTXGNRs)

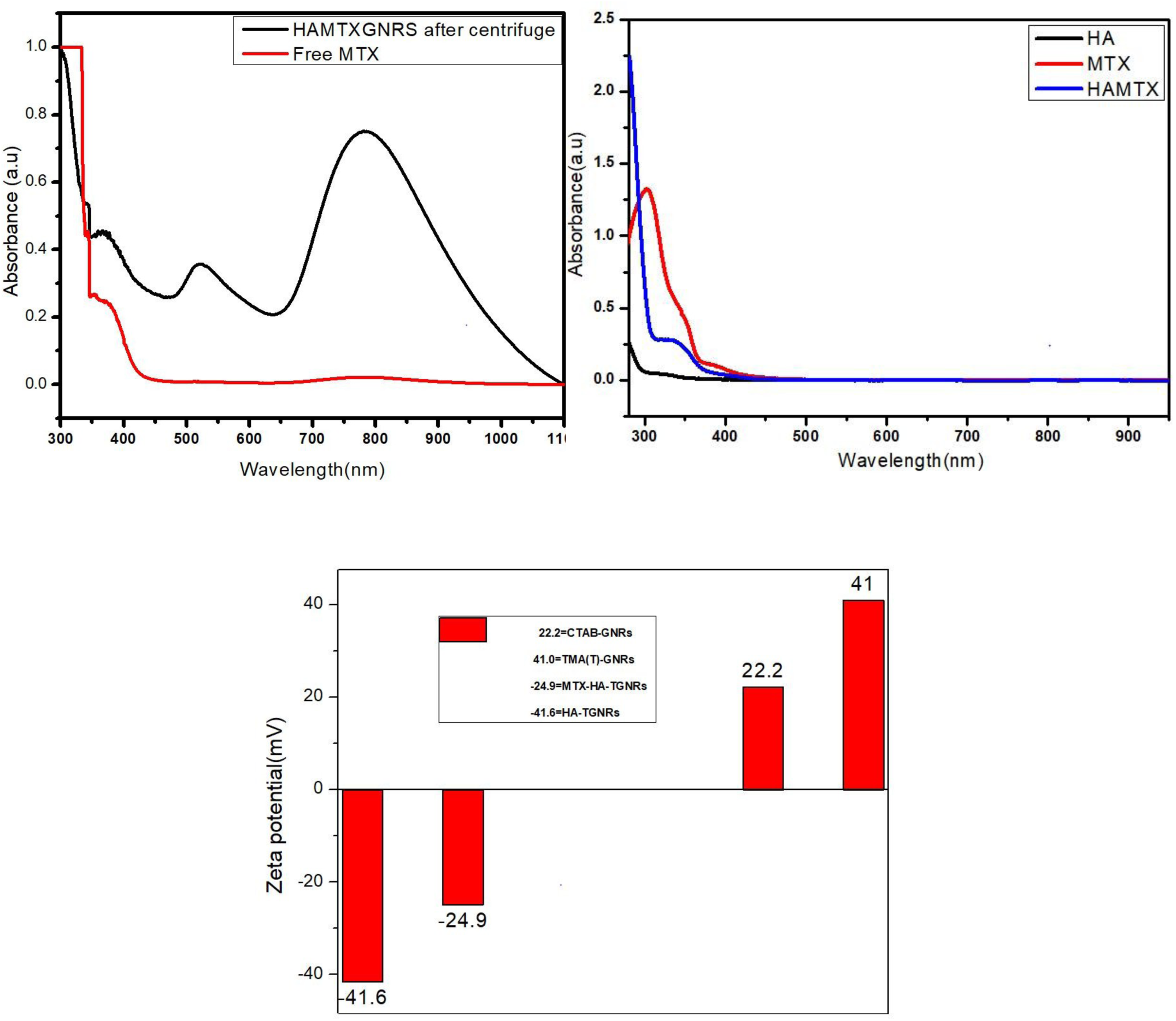

## (S3) Flu. Intensity of loaded amount of Rho. B on HAGNRs

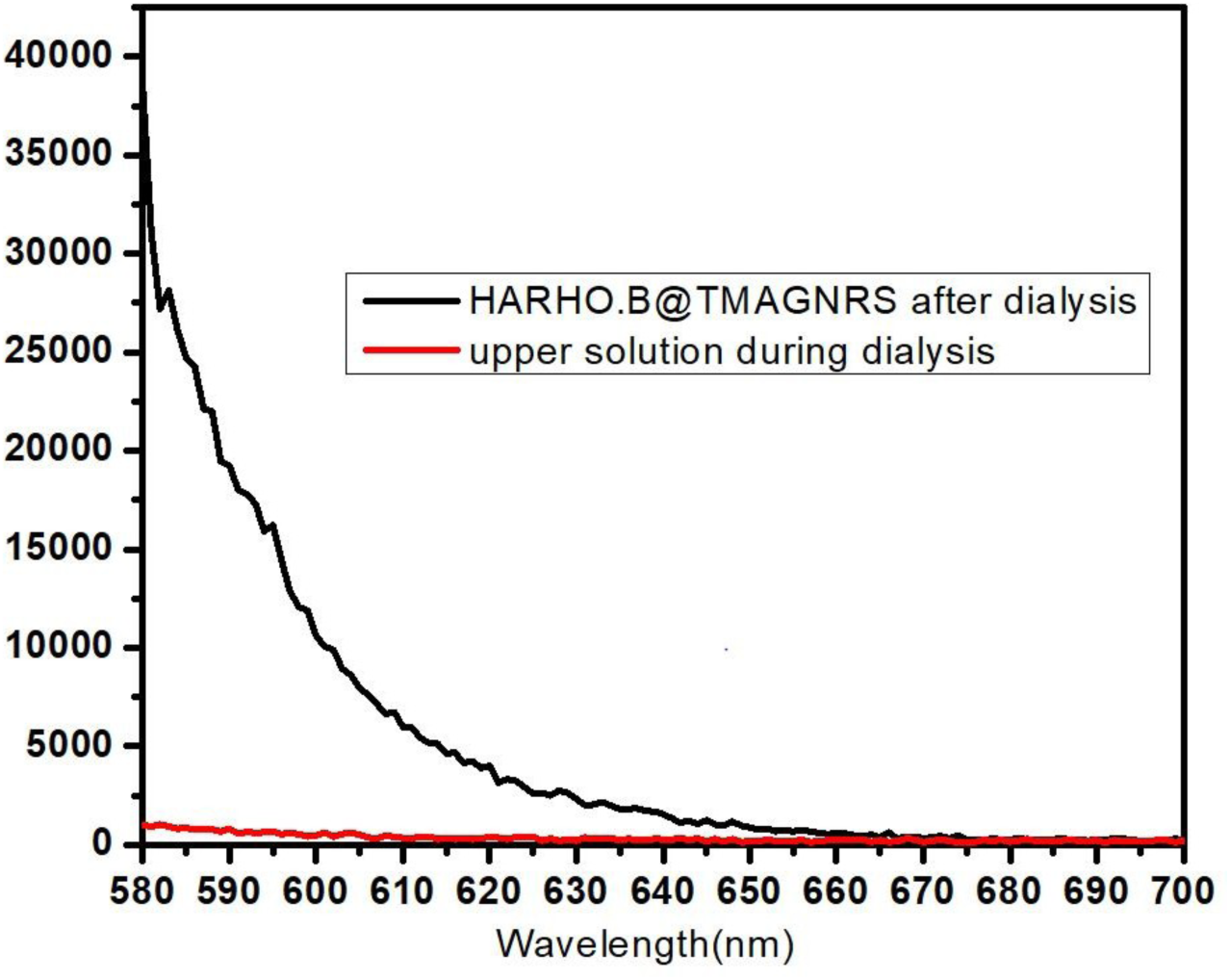

## (S4) Flu. Intensity of free Rho. B released from HARho.B (24h) at different Temperature in PBS

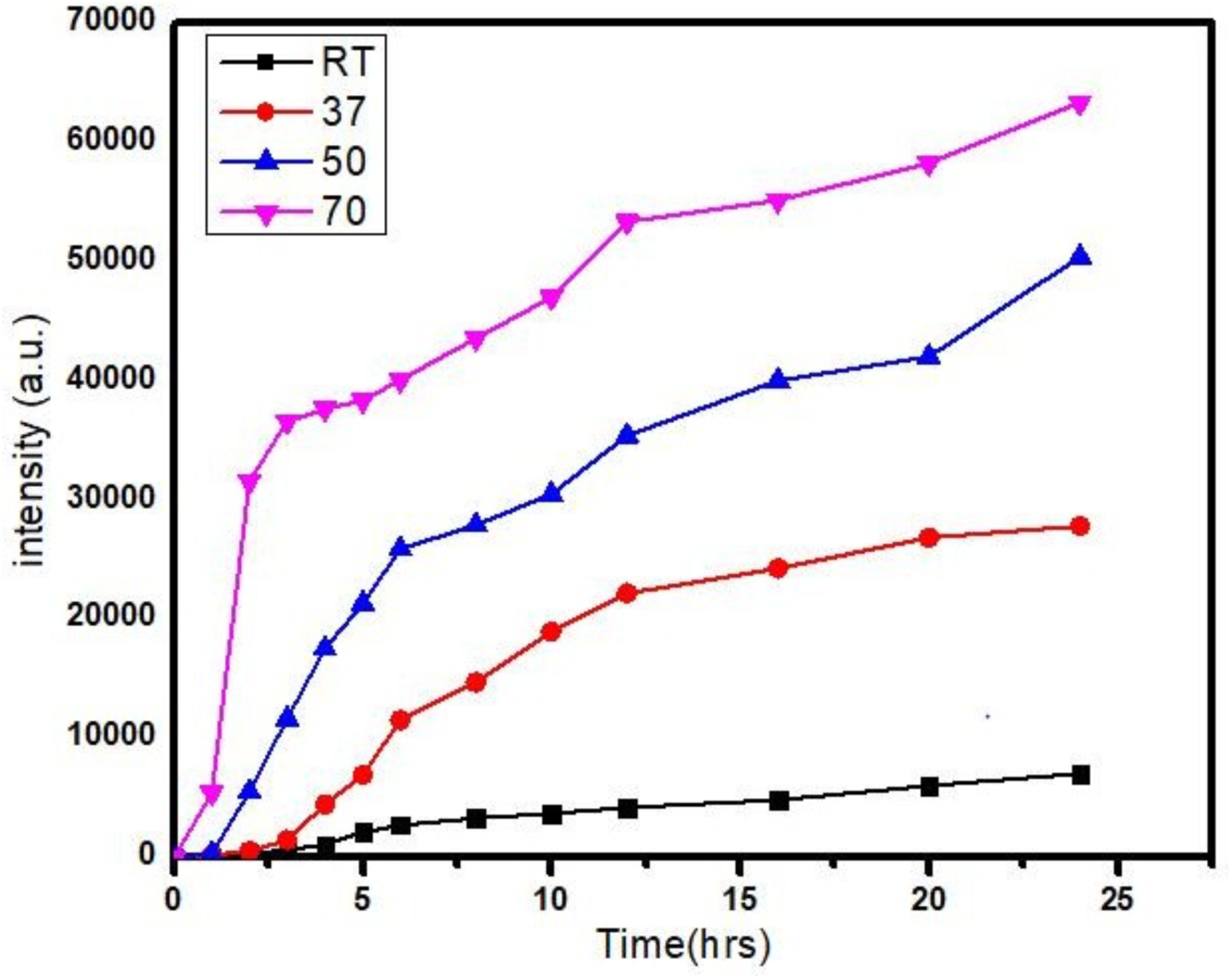

